# Sparse reduced-rank regression for exploratory visualization of paired multivariate datasets

**DOI:** 10.1101/302208

**Authors:** Dmitry Kobak, Yves Bernaerts, Marissa A. Weis, Federico Scala, Andreas Tolias, Philipp Berens

**Affiliations:** Institute for Ophthalmic Research, University of Tübingen, Germany; International Max Planck Research School for Intelligent Systems, Germany; Department of Neuroscience, Baylor College of Medicine, Houston, Texas, USA; Department of Computer Science, University of Tübingen, Germany

## Abstract

In genomics, transcriptomics, and related biological fields (collectively known as *omics*), it is common to work with *n* ≪ *p* datasets with the dimensionality much larger than the sample size. In recent years, combinations of experimental techniques began to yield multiple sets of features for the same set of biological replicates. One example is Patch-seq, a method combining single-cell RNA sequencing with electrophysiological recordings from the same cells. Here we present a framework based on sparse reduced-rank regression for obtaining an interpretable visualization of the relationship between the transcriptomic and the electrophysiological data. We use an elastic net regularization penalty that yields sparse solutions and allows for an efficient computational implementation. Using several publicly available Patch-seq datasets, we show that sparse reduced-rank regression outperforms both sparse full-rank regression and non-sparse reduced-rank regression in terms of predictive performance, and can outperform existing methods for sparse partial least squares and sparse canonical correlation analysis in terms of out-of-sample correlations. We introduce a *bibiplot* visualization in order to display the dominant factors determining the relationship between transcriptomic and electrophysiological properties of neurons. We believe that sparse reduced-rank regression can provide a valuable tool for the exploration and visualization of paired multivariate datasets, including Patch-seq.

## 1 Introduction

Since the days of Ramón y Cajal, neuroscientists have classified neurons into cell types, which are often considered the fundamental building blocks of neural circuits (Masland, 2004). Classically, these types have been defined based on their electrophysiology or anatomy, but due to the recent rise of single-cell transcriptomics, a definition of cell types based on genetics is becoming increasingly popular (Poulin et al., 2016). For example, single-cell RNA sequencing has been used to establish a census of neurons in the retina (Shekhar et al., 2016; Macosko et al., 2015), the cortex (Zeisel et al., 2015; Tasic et al., 2016, 2018), the whole brain (Saunders et al., 2018), and the entire nervous system (Zeisel et al., 2018) of mice. Despite this success, it has proven difficult to integrate the obtained cell type taxonomy based on the transcriptome with information about physiology and anatomy (Tripathy et al., 2017; Zeng and Sanes, 2017) and it remains unclear to what extent neural types are discrete or show continuous variation (Zeng and Sanes, 2017; Harris et al., 2018).

A recently developed technique called Patch-seq (Cadwell et al., 2016, 2017; Fuzik et al., 2016; Földy et al., 2016) allows to isolate and sequence RNA content of cells characterized electrophysiologically and/or morphologically (Figure 1a), opening the way to relate gene expression patterns to physiological characteristics on the single-cell level. Patch-seq experiments are laborious and low throughput, resulting in multimodal datasets with a particular statistical structure: a few dozen or hundreds of cells are characterized with expression levels of many thousands of genes as well as dozens of electrophysiological measurements (Figure 1a). Integrating and properly visualizing genetic and physiological information in this *n* ≪ *p* regime requires specialized statistical techniques that could isolate a subset of relevant genes and exploit information about the relationships within both data modalities to increase statistical power.

**Figure 1:**
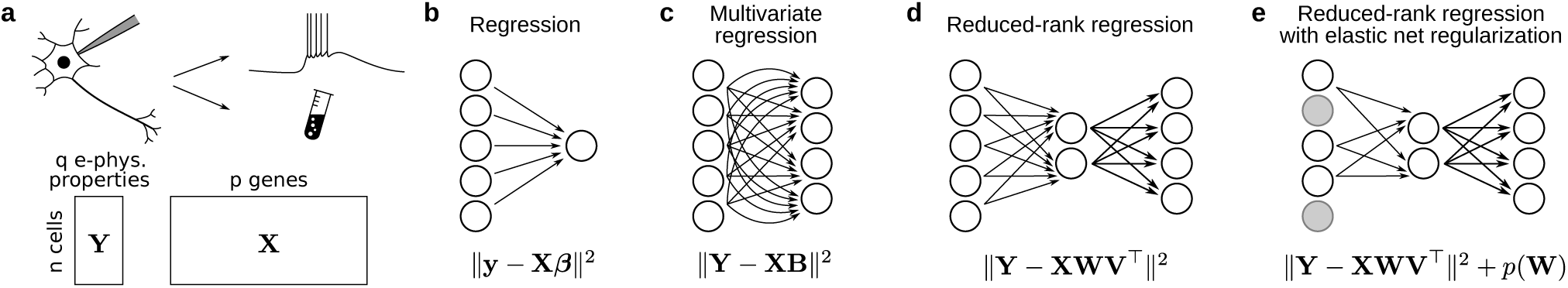
**a.** Schematic illustration of a Patch-seq experiment: electrophysiological activity is recorded by patch-clamping, followed by RNA extraction and sequencing. Below: data matrices after computational characterization of electrophysiological properties (**Y**) and estimation of gene counts (**X**). **b–e.** Schematic illustrations and loss functions for several regression methods. **b.** Simple regression. **c.** Multivariate regression. **d.** Reduced-rank regression. **e.** Regularized reduced-rank regression. Gray circles denote predictors that are left out of the sparse model.

Here we developed sparse reduced-rank regression based on the elastic net penalty to obtain an interpretable and intuitive visualization of the relationship between high-dimensional single-cell transcriptomes and electrophysiological information obtained using techniques like Patch-seq. We used four existing Patch-seq datasets (Fuzik et al., 2016; Cadwell et al., 2016; Scala et al., 2019, 2020) with sample sizes ranging from *n* = 44 to *n* = 1213 to demonstrate and validate our method (Table 1). Our sparse RRR method extends sparse RRR of Chen and Huang (2012) and, as we show, outperforms it on our data.

**Table 1:** Patch-seq datasets used in this study. All recordings were done in the mouse neocortex.

| Name | Citation | Cells ($n$) | Genes ($p$) | Features ($q$) | Description |
| --- | --- | --- | --- | --- | --- |
| M1 | Scala et al. (2020) | 1213 | 1000 | 16 | Motor cortex, all layers/types |
| L4 | Scala et al. (2019) | 102 | 1000 | 13 | Layer 4 *Sst* interneurons |
| L1 | Cadwell et al. (2016) | 44 | 3000 | 11 | Layer 1 interneurons |
| S1 | Fuzik et al. (2016) | 80 | 1384 | 80 | Layer 1/2 *Cck* interneurons |

Our code in Python is available at https://github.com/berenslab/patch-seq-rrr.

## 2 Results

### 2.1 Patch-seq data

A Patch-seq experiment yields two paired data matrices (Figure 1a): an *n × p* matrix **X** containing expression levels of *p* genes for each of the *n* cells, and an *n × q* matrix **Y** containing *q* electrophysiological properties of the same *n* cells. We assume that both matrices are centered, i.e. column means have been subtracted.

To illustrate the structure of such datasets and motivate the development of sparse RRR for exploratory visualization, we use principal component analysis (PCA) on the M1 dataset, one of the largest existing Patch-seq datasets (Scala et al., 2020). It contains *n* = 1213 neurons from the primary motor cortex of adult mice and spans all types of neurons, both excitatory and inhibitory (Table 1). Each cell was described by *q* = 16 electrophysiological properties and we used the *p* = 1000 most variable genes that were selected in the original publication. Note that there are over 40 thousand coding and non-coding genes in the mouse genome that were detected in at least one cell in this particular data set. It is, however, a common practice to select a smaller set of genes for downstream analysis (Luecken and Theis, 2019), as most detected genes have low average expression, low variance, and are likely not informative. We log-transformed all gene counts and standardized the columns of **X** and **Y** matrices (see Methods).

PCA in the transcriptomic space (Figure 2a) revealed that PC1, in this case, was an experimental artefact largely driven by the variability in the sequencing quality between cells (correlation between PC1 and the log number of detected genes was 0.90), whereas PC2 captured a biologically meaningful difference between the excitatory and the inhibitory cells. In contrast, PCA in the electrophysiological space (Figure 2b) separated major classes of neurons with different firing properties, such as *Pvalb*- (red), *Sst*- (orange), or *Vip*- (purple) expressing interneurons. Thus, there appears to be no direct relationship between the leading PCs of the two modalities. The aim of reduced-rank regression is to uncover such relationships.

**Figure 2:**
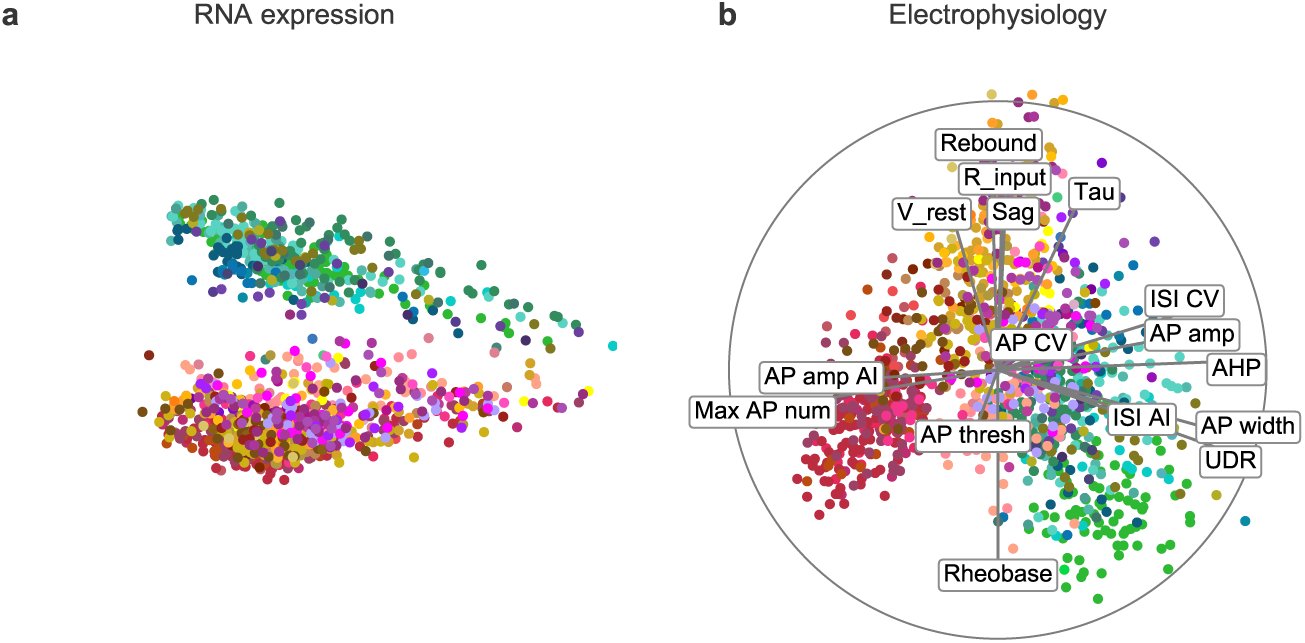
**a.** Principal component analysis (PCA) of the transcriptomic data in the M1 dataset (Scala et al., 2020). Color denotes transcriptomic type(cold colors: excitatory neurons; warm colors: inhibitory neurons). Both PCs were standardized. **b.** PCA biplot of the electrophysiological data in the same dataset. Grey lines show correlations of individual electrophysiological features with PC1 and PC2. The circle (*correlation circle*) shows maximal possible correlations. The relative scaling of the scatter plot and the lines/circle is arbitrary. The label positions were automatically adjusted by simulating spring repulsive forces between them until they stopped overlapping.

The visualisation in Figure 2b is known as *biplot* (Gabriel, 1971). Lines represent correlations between each electrophysiological property and PC1/PC2: the horizontal coordinate of each line’s tip shows the correlation with PC1 and the vertical coordinate shows the correlation with PC2. The circle, sometimes called *correlation circle*, shows the maximum attainable correlation. The scaling between the scatter plot and the lines/circle is arbitrary. Following Gabriel (1971), we standardize both PCs and scale the lines/circle by an arbitrary factor of 3 (so that most points in the scatter plot are contained within the circle). We do not show a biplot in the transcriptomic space (Figure 2a) because the PCA in the gene space is not sparse, making the biplot practically impossible to display and interpret as it would have to show all 1000 genes from **X**. This motivates the sparsity constraint that we impose on RRR.

### 2.2 Reduced-rank regression

To relate gene expression patterns to electrophysiological properties, one could use the transcriptomic data to predict any given electrophysiological property, e.g. action potential threshold. This is a *regression* problem: each gene is a predictor and the action potential threshold is the response variable (Figure 1b). To predict multiple electrophysiological properties at the same time, one can combine individual regressions into a *multivariate regression* problem where the response is a multivariate vector (Figure 1c). The loss function of multivariate linear regression (known as ordinary least squares, OLS) is

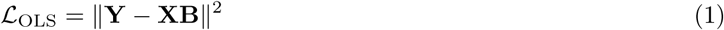

and its well-known solution is given by

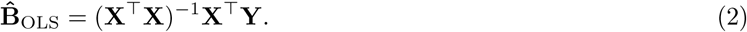

Here and below all matrix norms are Frobenius norms. The intercept is omitted because both **X** and **Y** are assumed to be centered.

Different electrophysiological properties tend to be strongly correlated and so one can construct a more parsimonious model where gene expression is predicting *r < q* latent factors that in turn predict all *q* electrophysiological properties together (Figure 1d). These latent factors form a bottleneck in the linear mapping and allow exploiting correlations between the predicted elecrophysiological properties to increase statistical power and decrease overfitting. This approach is called *reduced-rank regression* (RRR) (Izenman, 1975; Velu and Reinsel, 2013). Its loss function is

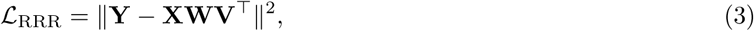

where **W** and **V** each have *r* ≤ min(*p, q*) columns. Without loss of generality it is convenient to require that **V**^⊤^**V** = **I**. The product **WV**^⊤^ forms the matrix of regression coefficients that has rank *r*.

This decomposition allows to interpret **W** as a mapping that transforms **X** into *r* latent variables and **V** as a mapping that transforms the latent variables into **Y** (Figure 1e). As a result, RRR can be viewed not only as a prediction method, but also as a dimensionality reduction method, allowing visualization and exploration of the paired dataset. Latent factors **XW** can be interpreted as capturing low-dimensional genetic variability that is predictive of electrophysiological variability, while **YV** can be interpreted as low-dimensional electrophysiological variability that can be predicted from the genetic variability.

RRR can be directly solved by applying singular value decomposition (SVD) to the results of multivariate regression. Indeed, the RRR loss can be decomposed into the OLS loss and the low-rank loss:

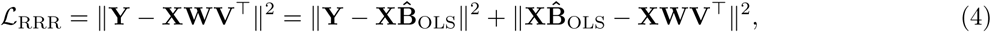

The first term corresponds to the variance of **Y** that is unexplainable by any linear model. The minimum of the second term can be obtained by computing the SVD of 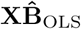. The right singular vectors corresponding to the *r* largest singular values give 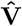, and 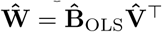.

Reduced-rank regression with the ridge penalty *λ* ∥**WV** ∥^2^ = *λ* ∥**W**∥ ^2^ has the same analytic solution, but 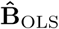 should be replaced with 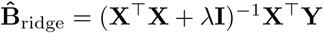.

### 2.3 Reduced-rank regression with elastic net penalty

As there are over 20 thousand genes in a mouse genome (with 1000–5000 typically retained for analysis) while the typical sample size *n* of a Patch-seq dataset is on the order of 100–1000, the regression problems discussed above are in the *n < p* regime and need to be regularized. Here we use elastic net regularization, which combines *ℓ*_1_ (lasso) and *ℓ*_2_ (ridge) penalties (Zou and Hastie, 2005). Elastic net enforces sparsity and performs feature selection: only a small subset of genes are selected into the model while all other genes get zero regression coefficients (Figure 1e). Our elastic net RRR extends a previously suggested sparse RRR (Chen and Huang, 2012) that used the lasso penalty on its own. The elastic net penalty has well-known advantages compared to the pure lasso penalty, e.g. it allows to select more than *n* predictors and can outperform lasso when predictors are strongly correlated (Zou and Hastie, 2005).

The loss function of our regularized RRR is:

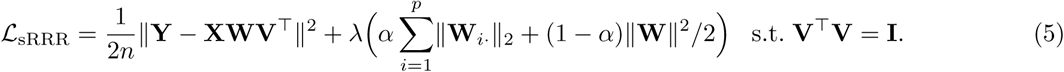

The *ℓ*_2_ penalty is only applied to the matrix **W** because **V** is constrained to have a fixed *ℓ*_2_ norm. Also, following Chen and Huang (2012) we chose not to apply *ℓ*_1_ penalty to **V** and impose sparsity constraint only on the gene selection (see Discussion). We used the same parametrization of the penalty as in the popular `glmnet` library (Friedman et al., 2010): *α* controls the trade-off between the lasso (*α* = 1) and the ridge (*α* = 0) while *λ* controls the overall regularization strength. Following Chen and Huang (2012), the lasso penalty term 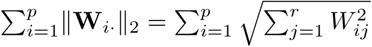 computes the sum of *ℓ*_2_ norms of each row of **W**. This is known as *group lasso* (Yuan and Lin, 2006), because it is the *ℓ*_1_ norm of the vector of row *ℓ*_2_ norms; it encourages the entire rows of **W**, and not just its individual elements, to be zeroed out, corresponding to some of the genes being left out of the model entirely. See Discussion about this choice.

This optimization problem is biconvex and can be solved with an iterative alternating approach: in turn, we fix **V** and find the optimal **W**_opt_ and then fix **W** and find the optimal **V**_opt_ until convergence. For a fixed **V**, the least-squares term can be re-written as

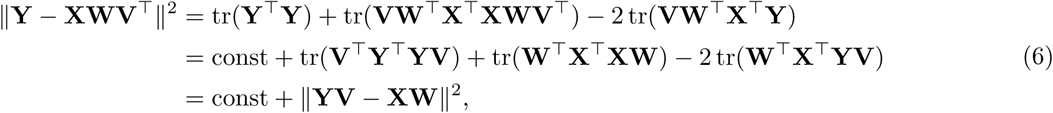

meaning that for a fixed **V**, the loss is equivalent to

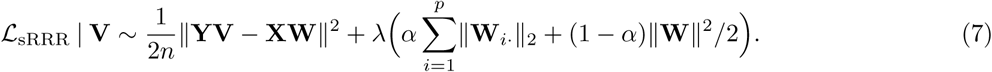

This is the loss of multivariate elastic net regression of **YV** on **X**, and so the optimal **W**_opt_ can be obtained using the `glmnet` library (Friedman et al., 2010) (using `family=“mgaussian”` option for row-wise lasso penalty) which has readily available interfaces for Matlab, Python, and R.

For a fixed **W**, the loss does not depend on the penalty terms and the least-squares term can be written as

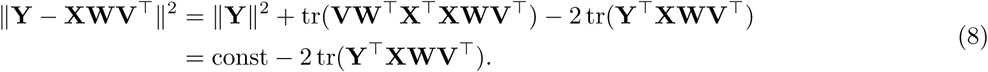

This is an example of the orthogonal Procrustes problem (Gower and Dijksterhuis, 2004). Maximizing tr(**Y**^⊤^**XWV**^⊤^) is achieved by the thin SVD of **Y**^⊤^**XW**. If the *r* left and right singular vectors are stacked in columns of **L** and **R** respectively (we order them by singular values, in decreasing order), then **V**_opt_ = **LR**^⊤^. We provide a short proof in the Appendix.

Given that the loss function is biconvex but possibly not jointly convex in **V** and **W**, it can be important to choose a reasonable initialization. We initialized **V** by the *r* leading right singular vectors of **X**^⊤^**Y** and found this strategy to work well.

### 2.4 Relaxed elastic net

It has been argued that elastic net or even the lasso penalty on its own can lead to an over-shrinkage with non-zero coefficients shrinking too much (Zou and Hastie, 2005). There have been several suggestions in the literature on how to mitigate this effect (Efron et al., 2004; Zou and Hastie, 2005; Meinshausen, 2007). *Relaxed lasso* (Meinshausen, 2007) performs lasso (setting *α* = 1 and *λ* = *λ*_1_) and then, using only the terms with non-zero coefficients, performs another lasso with a different penalty (*α* = 1, *λ* = *λ*_2_; usually *λ*_2_ *< λ*_1_). If *λ*_2_ = 0, then this has also been called *LARS-OLS hybrid* (Efron et al., 2004).

Similar two-stage procedures for the elastic net penalty are not as established. We obtained a strong improvement in predictive performance if — after RRR with elastic net penalty with coefficients *λ* and *α* — we take the genes with non-zero coefficients and run RRR again using *α* = 0 (i.e. pure ridge) and the same value of *λ*. This procedure does not introduce any additional tuning parameters but substantially outperforms pure elastic net RRR on our data, as we show below. We called it *relaxed elastic net*, following the relaxed lasso terminology (Meinshausen, 2007). The solution of the first round of sparse RRR we call *naïve*, following Zou and Hastie (2005).

A similar approach was used by De Mol et al. (2009) who performed elastic net using *λ* = *λ*_1_ and some small fixed value of *α* = *α*_1_, selected all genes with non-zero coefficients, and did pure ridge (*α* = 0) regression with *λ* = *λ*_2_ on this gene subset. This approach also has two hyperparameters that need to be selected using cross-validation, but requires a manual choice of *α*_1_ for the first elastic net. If *α* is also treated as an adjustable hyperparameter, then it becomes a more flexible generalization of our approach with three hyperparameters.

### 2.5 Cross-validation

We used cross-validation (CV) to select the values of *r, λ*, and *α* that maximize the predictive performance of the sparse RRR model. The cross-validation estimates of *R*^2^ are shown in Figure 3 for the L1, L4, and M1 datasets. We used 10 times repeated 11-fold CV for the L1 data (*n* = 44), 10 times repeated 10-fold CV for the L4 data (*n* = 102), and non-repeated 10-fold CV for the M1 data (*n* = 1321). See Methods for the pre-processing details. The test-set *R*^2^ for each CV fold was computed as

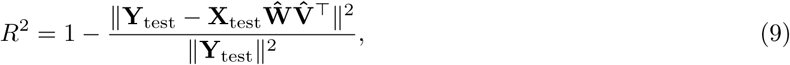

where **X**_test_ and **Y**_test_ were centered using the corresponding training-set means. We averaged the resulting *R*^2^ across all folds and repetitions.

**Figure 3:**
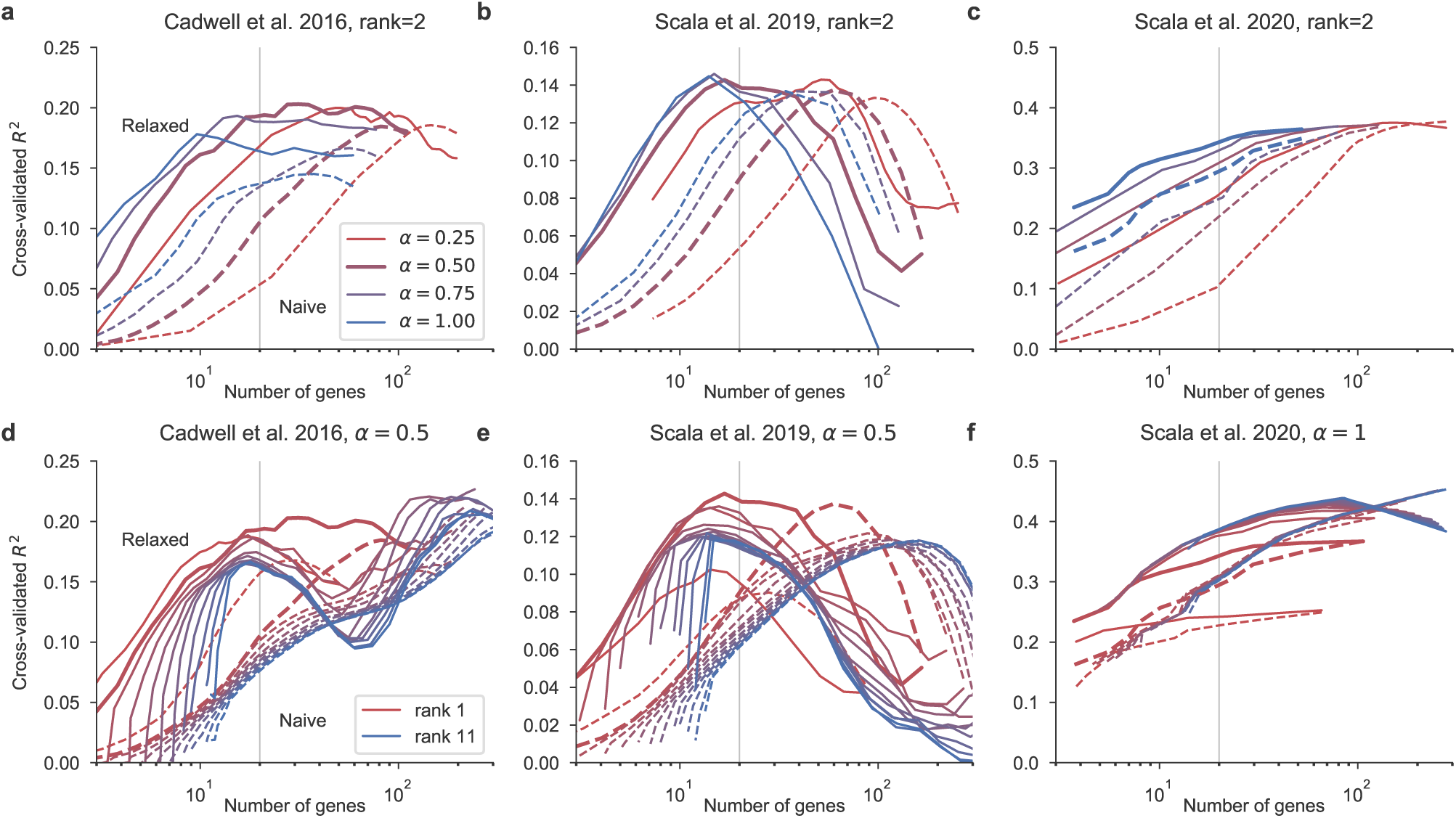
**a.** Cross-validation performance of sparse RRR with *r* = 2 on the L1 dataset, depending on *α* (color-coded, see legend) and *λ*. Horizontal axis shows the average number of selected genes obtained for each *λ*. Dashed lines: naive sparse RRR. Solid lines: relaxed sparse RRR. The vertical line at 20 selected genes indicates our parameter choice. Thick lines highlight *α* = 0.5. The standard deviation over all CV folds and repetitions at *α* = 0.5 and *λ* value yielding ∼20 genes was 0.05 for the naive and 0.12 for the relaxed estimator (note that each fold had only 4 samples in the test set). **b.** The same for the L4 dataset. The standard deviation over all CV folds and repetitions at *α* = 0.5 and *λ* value yielding ∼20 genes was 0.04 for the naive and 0.07 for the relaxed estimator (here each fold had only ∼10 samples in the test set). **c.** The same for the M1 dataset. Here thick lines highlight *α* = 1. The standard deviation over all CV folds and repetitions at *α* = 1 and *λ* value yielding ∼20 genes was 0.01 for the naive and for the relaxed estimators (here each fold had ∼120 samples in the test set). **d.** Cross-validation performance with *α* = 0.5 depending on the rank (color-coded, see legend) on the L1 dataset. Thick lines highlight *r* = 2. **e.** The same for the L4 dataset. **f.** The same for the M1 dataset (here using *α* = 1).

When using rank *r* = 2 (Figure 3a–c), we found that *α* = 0.5 outperformed *α* = 1 on the L1 dataset, suggesting that adding an additional ridge penalty to the sparse RRR model of Chen and Huang (2012) can be helpful. At the same time, *α* = 0.5 and *α* = 1 performed equally well on the L4 dataset, while *α* = 1 outperformed other values on the M1 dataset. Overall, the differences in predictive performance in the *α* ∈ [0.5, 1] range were moderate. For the downstream analysis, we used *α* = 0.5 for the L4 and L1 datasets, and *α* = 1 for the M1 dataset. We recommend *α* = 0.5 as a default setting.

The optimal *λ* corresponded to ∼30 selected genes for the L1 dataset, ∼15 selected genes for the L4 dataset, and ∼100 selected genes for the M1 dataset (Figure 3a–c), but the performance was comparably good in the range of ∼10–100 genes. For the downstream analysis, we always chose the value of *λ* yielding 20 selected genes. Selecting many more genes than that would make visualisation difficult (see below).

The optimal value of rank was *r* = 2 for the L1 and L4 datasets and *r* ≈ 10 for the M1 dataset (Figure 3d–f). Lower ranks had worse performance due to under-fitting, whereas higher ranks led to a drop in performance due to over-fitting. Note that the full rank (*r* = 11, *r* = 13, and *r* = 16 for the L1, L4, and M1 datasets respectively) corresponds to the standard multivariate elastic net regression. We verified that our algorithm yields the same solution as `glmnet` does on its own. The much better performance of *r* = 2 compared to the full rank on the L1 and L4 datasets shows that *r* can act as a regularization parameter, making sparse reduced-rank regression outperform sparse full-rank regression. At the same time, for the M1 dataset, there was almost no difference in performance for any *r* ≥ 5 and the full-rank model performed almost as well, suggesting little overfitting due to the larger sample size. Still, even in this case, the RRR framework allows to order individual components by their importance (explained variance) and to make low-dimensional visualizations (see below).

Finally, in both datasets the relaxed version of sparse RRR strongly outperformed the naive version, at least in the range of 10–50 selected genes, which is the range needed for visualizations (see below). If a high number of genes was selected into the model, the relaxed version performed worse than the naive version, suggesting that the second stage of our relaxed approach was overfitting. For the L1 dataset with the smallest sample size we observed non-monotonic dependency of the relaxed performance on the number of genes (Figure 3d), suggesting that the relaxed estimator can occupy different positions on the bias/variance trade-off depending on the *λ*. However, for the low number of selected genes the relaxed version had superior performance across all datasets, all ranks, and all values of *α*.

Note that sparse RRR of Chen and Huang (2012) corresponds to the naive version with *α* = 1. In the regime when the model selects a few dozen genes, it was strongly outperformed by our relaxed sparse RRR estimator.

### 2.6 Bibiplot visualisation

We applied our sparse RRR approach with *r* = 2 and *λ* chosen to yield 20 selected genes to the L1, L4, and M1 datasets (used *α* values: 0.5, 0.5, and 1.0 respectively). For each of the datasets, we visualized the results with a pair of biplots, a graphical technique that we suggest to call a *bibiplot*.

To construct a biplot in the transcriptomic space, we use the bottleneck representation **XW** for the scatter plot, and show lines for all genes that are selected by the model (even though other genes can also have non-zero correlations with **XW**). The biplot in the electrophysiological space is constructed using **YV** and shows all available electrophysiological properties. If *R*^2^ of the model is high, then the two scatter plots will be similar to each other. Comparing the directions of variables between the two biplots can suggest which electrophysiological variables are associated with which genes.

The L1 dataset encompasses two types of interneurons from layer 1 of mouse cortex: neurogliaform cells (NGC) and single bouqet cells (SBC). Accordingly, the first RRR component captured the difference between the two cell types (Figure 4a,d). The second RRR component had only one gene strongly associated with it (Figure 4a) and contributed only a very small increase in cross-validated *R*^2^, as one can see comparing the cross-validation curves for *r* = 1 and *r* = 2 (Figure 3d). We conclude that the second RRR component in this dataset is only weakly detectable.

**Figure 4:**
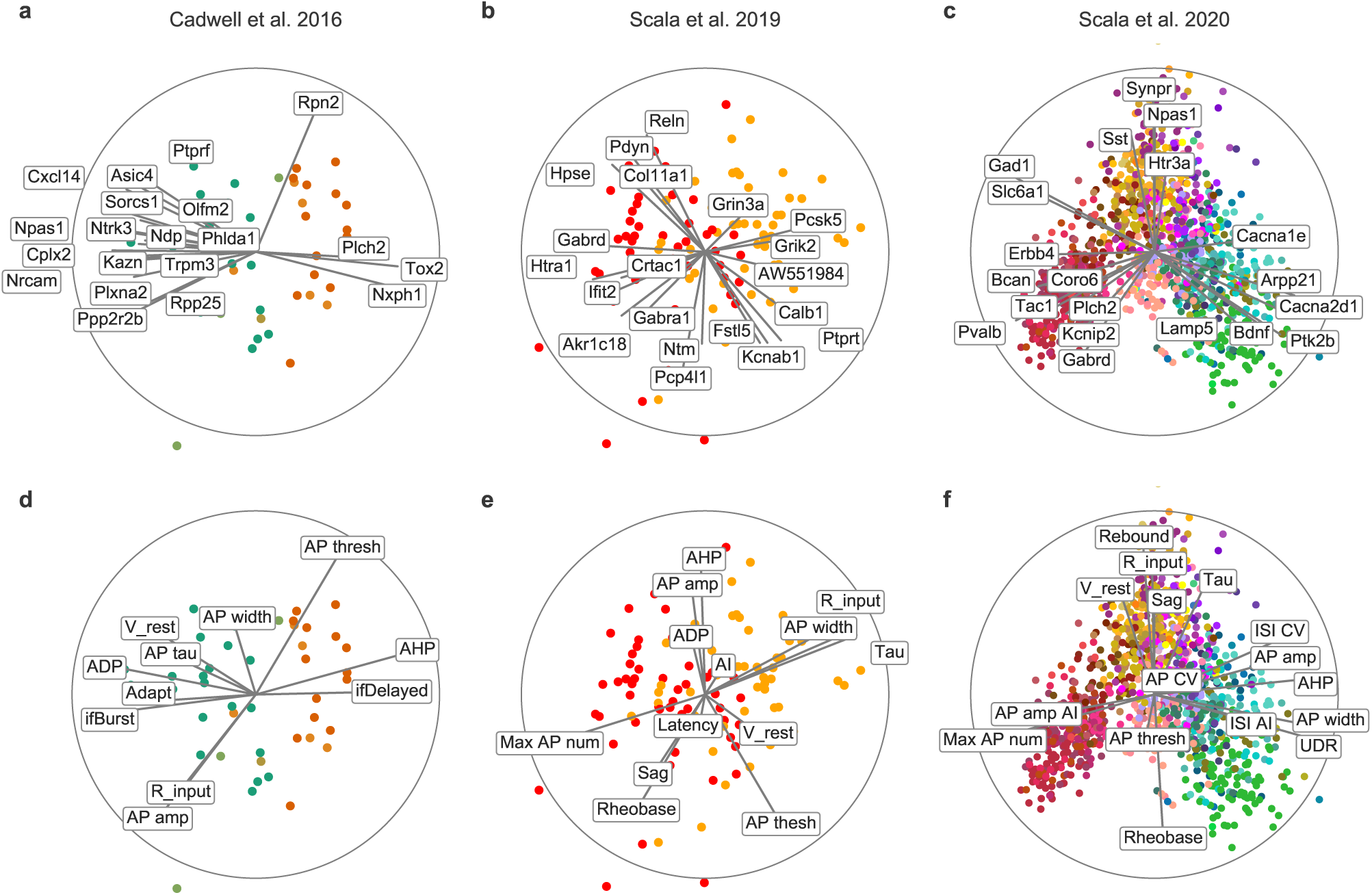
**a.** Sparse RRR biplot of the transcriptomic space in the L1 dataset. Color codes cell type (orange: neurogliaform cells, NGC; green: single bouqet cells, SBC). Only 20 genes selected by the model are shown. See Figure 2b for the details of biplot visualization. As there, label positions were automatically adjusted to prevent overlap. **b.** The same for the L4 dataset. Color denotes cortical area (orange: visual cortex; red: somatosensory cortex). **c.** The same for the M1 dataset. Color denotes transcriptomic type (see Figure 2). **d–f.** The same in the electrophysiological spaces.

In the L4 dataset (Figure 4b,e), the most salient feature in the bibiplot is the separation between the cells recorded in the visual and the somatosensory cortices. The selected genes here are pointing in all directions, and indeed the second component contributed a substantial increase in *R*^2^ (Figure 3e). This suggests that both components are biologically meaningful. See Scala et al. (2019) for a more in-depth analysis using sparse RRR.

The M1 dataset was much larger than the other two and included a much more diverse selection of neuron types. As a result, the *R*^2^ values were substantially higher and the model needed to use rank *r* ≥ 5 to reach its optimal performance. Here we nevertheless used *r* = 2 because it allows the same kind of visualization as for the other datasets (Figure 4c,f). See Scala et al. (2020) for a more in-depth analysis using sparse RRR with rank *r* = 5. Two-dimensional bibiplot separated major classes of neurons, such as *Pvalb, Sst, Vip*, and *Lamp5* expressing interneurons (red/orange/purple/salmon), and excitatory cells (green). It is worth noting that the RRR biplot in the electrophysiological space (Figure 4f) was very similar to the PCA biplot (Figure 2b). This indicates that the sparse RRR model explained the dominant modes of variation among the dependent variables. We observed the same in other datasets analyzed here.

### 2.7 Comparison to sparse CCA and PLS

Reduced-rank regression does not directly aim to maximize the correlation between **Xw** and **Yv**, where **w** and **v** are corresponding columns of **W** and **V**, even though high correlation is needed to achieve high *R*^2^. Nevertheless, one can ask what is the cross-validated estimate of this correlation in each pair of RRR components. We used the same cross-validation scheme to measure these out-of-sample correlations (Figure 5)^1^. With the hyper-parameters used above, the correlations in the L1 dataset were 0.70 for component 1 and 0.37 for component 2 (Figure 3a). In the L4 dataset, they were 0.65 and 0.54, respectively (Figure 3b). In the M1 dataset, they were 0.89 and 0.73 (Figure 1c).

**Figure 5:**
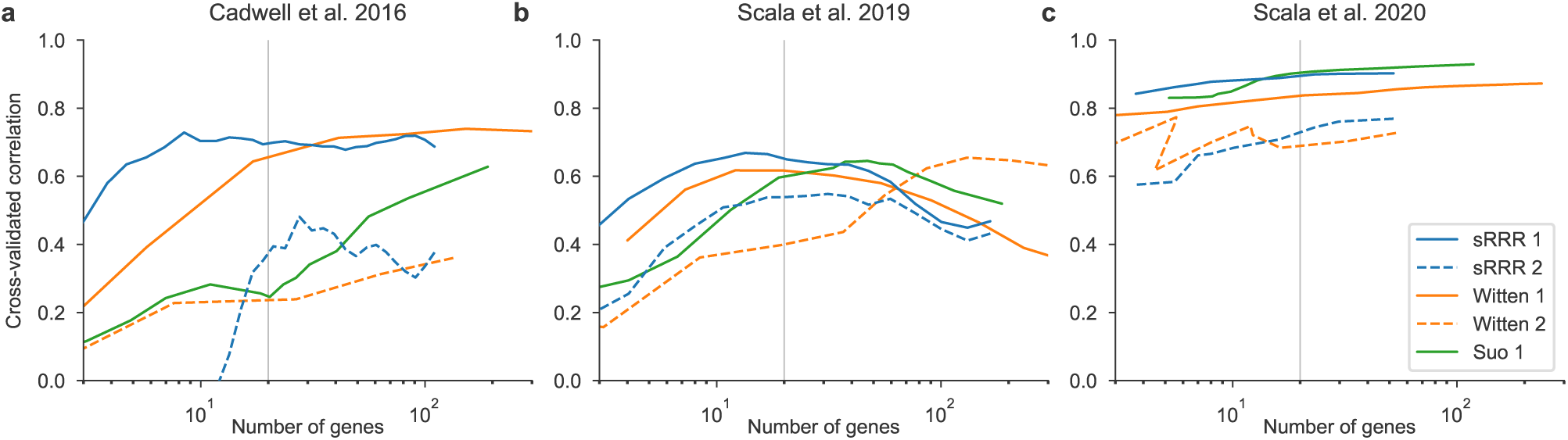
**a.** Cross-validation estimates of correlations between the transcriptomic and the electrophyiological RRR components with *r* = 2 in the L1 dataset, depending on *λ*. Horizontal axis shows the average number of selected genes obtained for each *λ*. Here we used *α* = 0.5. Solid blue line: RRR component 1. Dashed blue line: RRR component 2. Solid and dashed orange lines: sparse CCA method of Witten et al. (2009), components 1 and 2. Green line: sparse CCA method of Suo et al. (2017), component 1. **b.** The same for the L4 dataset. **c.** The same for the M1 dataset.

A statistical method that directly maximizes correlation between **Xw** and **Yv** is called canonical correlation analysis (CCA). A number of different methods for sparse CCA have been suggested in the last decade (Waaijenborg et al., 2008; Wiesel et al., 2008; Parkhomenko et al., 2009; Witten et al., 2009; Witten and Tibshirani, 2009; Lykou and Whittaker, 2010; Hardoon and Shawe-Taylor, 2011; Chen et al., 2012b; Chu et al., 2013; Wilms and Croux, 2015; Gao et al., 2017; Suo et al., 2017), of which the sparse CCA of Witten et al. (2009) is arguably the most well-known (judging by the number of citations in Google Scholar). We reimplemented the algorithm of Witten et al. in Python and used the cross-validation procedure described above to measure its out-of-sample performance (Figure 5, blue lines). We found that it performed noticeably worse than our sparse RRR: correlations for all three datasets and both components (the 1st and the 2nd) were similar or lower than with sparse RRR, at least in the regime of 10–50 selected genes. We also implemented the recently suggested sparse CCA algorithm of Suo et al. (2017) that directly builds up on the approach of Witten et al. We found that the sparse CCA of Suo et al. performed worse than the sparse CCA of Witten et al. in the L1 dataset but outperformed it in the M1 dataset, where its performance was similar to our sparse RRR (Figure 5, green lines).

It remains beyond the scope of this paper to investigate why our sparse RRR is competitive with or even outperforms these sparse CCA methods in terms of out-of-sample correlations. A likely explanation is that these sparse CCA methods use regularization approaches that are suboptimal for our datasets. To understand this under-/over-regularization, note that RRR maximizes explained variance in **Y**, i.e. correlation between **Xw** and **Yv**, times the standard deviation of **Yv**. Another related method is called partial least squares (PLS): it maximizes the covariance between **Xw** and **Yv**, i.e. correlation, times the standard deviation of **Yv**, times the standard deviation of **Xw**. In some sense, both RRR and PLS can be seen as particular regularized versions of CCA, because they bias **w** and **v** towards the high-variance directions in **X** and **Y**, somewhat similar to the ridge penalty. The method of Witten et al. maximizes covariance (and so could in fact have been called sparse PLS and not sparse CCA), which might provide too strong *ℓ*_2_ regularization. The method of Suo et al. maximizes correlation and does not use any ridge penalty, which might cause too weak *ℓ*_2_ regularization.

### 2.8 Gene selection stability

Instability is a general feature of all sparse models, especially when *n* ≪ *p* (Xu et al., 2011). We used bootstrapping to estimate gene selection stability in our datasets. On each of the 100 iterations, we drew a bootstrap sample of *n* cells with repetitions and fit the sparse RRR model with the same parameters as above. This allows to measure how often each gene is selected into the model.

As expected, we found that the larger the sample size the more stable the gene selection was. In the L1 dataset (*n* = 44), the most reliably selected gene was selected only 67% of times. In the L4 dataset (*n* = 102), 89% of times. In the M1 dataset (*n* = 1213), there were 10 genes that were selected over 90% of times, with several genes getting into the model on every bootstrap iteration.

We also found that elastic net with *α* = 0.5 led to a more stable model than the pure lasso with *α* = 1 (with *λ* values appropriately adjusted to select 20 genes). We quantified the overall gene selection stability by computing the mean and the standard deviation of the bootstrap selection fraction across the top 20 most often selected genes. This average stability was 0.24 *±* 0.10 with *α* = 1.0 vs. 0.34 *±* 0.12 with *α* = 0.5 in the L1 dataset; 0.48 *±* 0.12 vs. 0.55 *±* 0.19 in the L4 dataset; and 0.80 *±* 0.20 vs. 0.85 *±* 0.19 in the M1 dataset. The difference was not large, but consistently observed in each dataset. See Discussion for more considerations about model stability.

### 2.9 Preprocessing choices

All of the analysis shown above was done after selecting 1000–3000 most variable genes and standardizing the predictors. Putting these two preprocessing steps inside the cross-validation loop yielded practically the same results (Figure 6a–c; magenta lines).

**Figure 6:**
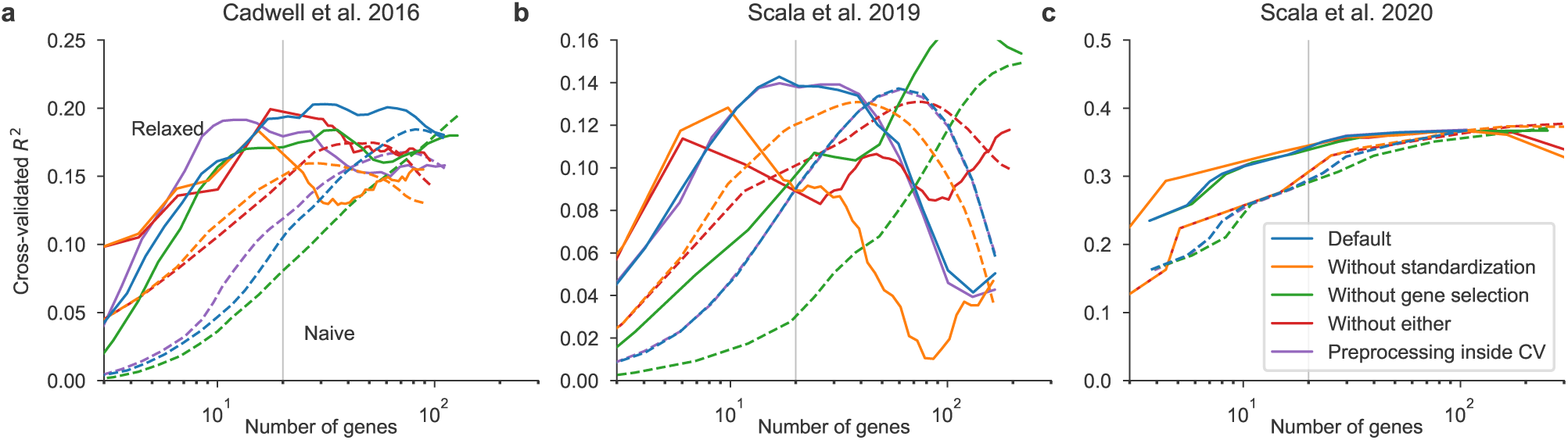
**a.** Cross-validated *R*^2^ in the L1 dataset with *r* = 2, *α* = 0.5, and different values of *λ*. Solid lines: relaxed version, dashed lines: naive version. Colors code different preprocessing choices. Blue: default approach used in other figures. Orange: without standardizing the columns of **X**. Green: without selecting most variable columns of **X**. Red: without either. Purple: gene selection and standardization done within the CV loop on each training set separately (and applied to the test set). **b.** The same for the L4 dataset. **c.** The same for the M1 dataset.

Omitting the gene selection step and performing sparse RRR directly on all detected genes (this number can exceed 40000, counting both coding and non-coding genes) and/or omitting the standardization step led to lower cross-validated *R*^2^ values in the L1 and L4 datasets but to exactly the same performance in the M1 dataset (Figure 6). This suggests that feature selection and standardization can be useful heuristics when the sample is low, but are not needed for larger sample sizes.

### 2.10 Sparse RRR with *r* ≠ 2

For the L1 and L4 datasets, cross-validation suggested *r* = 2 as the optimal rank, conveniently allowing us to use two-dimensional scatter plots for visualization. For the M1 dataset we used *r* = 2 for visualization, despite cross-validation suggesting that a higher rank. In this case, one can show several biplots for different pairs of components, or alternatively perform separate RRR analyses on the subsets of the data. We refer to our parallel publication describing the M1 dataset for further analysis (Scala et al., 2020).

We observed an opposite case when we applied sparse RRR to the S1 dataset with *n* = 80 inhibitory (all *Cck* from layers 1/2) and excitatory neurons from mouse somatosensory cortex (Fuzik et al., 2016). The first RRR component strongly separated excitatory and inhibitory neurons (Figure 7), which is not surprising given the large differences in gene expression and in firing patterns between these two classes of neurons. However, subsequent RRR components did not carry much signal in this dataset. The RRR model with *r* = 1 and *α* = 0.5 outperformed the model with *r* = 2 for low number of selected genes (Figure 7a), while the correlation in the 2nd component pair was close to zero (Figure 7b). This suggests effectively a one-dimensional shared subspace in this dataset.

**Figure 7:**
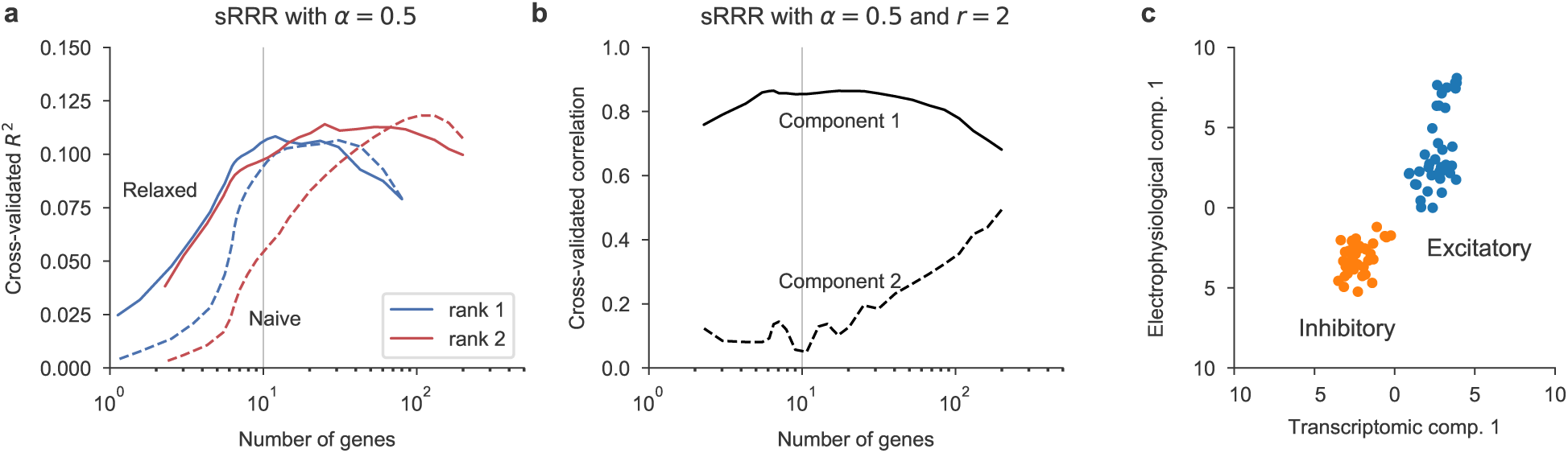
**a.** Cross-validation estimates of *R*^2^ in sparse RRR with *r* = 1 and *r* = 2 (*α* = 0.5) in the S1 dataset. Horizontal axis shows the average number of selected genes obtained for each *λ*. **b.** Cross-validation estimates of correlations between the transcriptomic and the electrophyiological RRR components with *r* = 2 and *α* = 0.5. **c.** The RRR component using *r* = 1 and *α* = 0.5 in the transcriptomic space (horizontal axis) and in the electrophysiological space (vertical axis). The value of *λ* was chosen to yield 10 selected genes.

## 3 Discussion

We proposed sparse reduced-rank regression (sRRR) as a tool for interpretable data exploration and visualization of Patch-seq recordings as an example of paired multivariate datasets in general. It allows to visualize the variability across cells in transcriptomic and electrophysiological modalities in a consistent way, and to find a sparse set of genes explaining electrophysiological variability. We used cross-validation to tune the hyper-parameters and to estimate the out-of-sample performance of the model.

### 3.1 Comparison to other regression methods

Our method directly builds up on the sparse RRR of Chen and Huang (2012) who added the lasso penalty to the RRR loss function. We extended this approach by using an elastic net penalty that combines lasso and ridge regularization. This adds flexibility to the method and indeed we showed that in some cases non-zero ridge penalty is beneficial for the predictive performance (Figure 3) and for model stability. In addition, we introduced the relaxed elastic net approach to mitigate the over-shrinkage bias associated with naive elastic net or naive lasso solutions (Efron et al., 2004; Zou and Hastie, 2005; Meinshausen, 2007; De Mol et al., 2009). The sparse RRR method of Chen and Huang (2012) corresponds to our naive RRR with *α* = 1, which in our experiments performed much worse than the relaxed version (Figure 3).

Furthermore, on some of the datasets sparse reduced-rank regression outperformed sparse full-rank regression (which is directly available e.g. in the popular `glmnet` library), suggesting that reduced-rank constraint is not only useful for visualisation but can also provide additional regularization and reduces overfitting.

Elastic net regularization has two parameters, *α* and *λ*, and cross-validation sometimes indicated that *α* can be varied in some range without affecting the model performance (Figure 3). This allows the researcher to control the trade-off between a sparser solution and a more comprehensive gene selection. If there is a set of genes that are highly correlated among each other, then large *α* will tend to select only one of them, whereas small *α* will tend to assign similar weights to all of them. We recommend using *α* = 0.5 as a reasonable default compromise.

We followed Friedman et al. (2010), Chen and Huang (2012) and others in imposing sparsity only in the predictor space: the lasso penalty is applied only to **W** but not to **V** and so feature selection only happens on the columns of **X** but not of **Y**. This arguably makes sense for the Patch-seq data considered in this manuscript but may be different for other kinds of datasets. See Chen et al. (2012a) for a discussion of sparse reduced-rank regression with sparsity in both **X** and **Y**.

### 3.2 Comparison to other dimensionality reduction methods

Reduced rank-regression is closely related to two other classical dimensionality reduction methods analyzing two paired data matrices (also called *two-view* data): canonical correlation analysis (CCA) and partial least squares (PLS). They can be understood as looking for projections with maximal correlation (CCA) or maximal covariance (PLS) between **X** and **Y**, whereas RRR looks for projections with maximal explained variance in **Y**. In recent years, multiple approaches to sparse CCA (Waaijenborg et al., 2008; Wiesel et al., 2008; Parkhomenko et al., 2009; Witten et al., 2009; Witten and Tibshirani, 2009; Lykou and Whittaker, 2010; Hardoon and Shawe-Taylor, 2011; Chen et al., 2012b; Chu et al., 2013; Wilms and Croux, 2015; Gao et al., 2017; Suo et al., 2017) and sparse PLS (Lê Cao et al., 2008, 2011; Chun and Keleş, 2010) have been suggested in the literature. Here, we chose sparse RRR at the core of our framework, because for the Patch-seq data it seems more meaningful to predict electrophyiological properties from transcriptomic information instead of treating them symmetrically, as genes give rise to physiological function. In addition, sparse RRR allows a mathematically simple formulation for rank *r >* 1 (using group lasso), whereas all sparse PLS/CCA methods cited above are iterative: after extracting the *k*-th component, matrices **X** and **Y** are deflated and the algorithm is repeated to extract the (*k* + 1)-th component, resulting in a cumbersome procedure that is often difficult to analyze mathematically.

That said, by comparing our sparse RRR method with sparse PLS/CCA methods of Witten et al. (2009) and Suo et al. (2017), we unexpectedly found that our method is competitive as a CCA variant. This suggests that there is room for new sparse CCA methods that would be more appropriate at least for the kind of datasets studied here.

Sparse (and non-sparse) CCA and PLS have been applied to biological datasets in order to integrate multiomics data (Lê Cao et al., 2008, 2009; González et al., 2008, 2009, 2012), with the recent mixOmics package for R providing a convenient implementation for several of these methods (Rohart et al., 2017). We believe that our sparse RRR can be a useful addition to this array of multi-omics statistical techniques.

This series of multi-omics papers always used two separate plots for visualizing the CCA/PLS results within each modality: a *sample plot*, also called a *units plot* (in our case this would be a scatter plot of Patch-seq cells), and a *variable plot*, also called a *correlation circle plot* (in our case this would be a scatter plot of selected genes or electrophysiological properties, together with the correlation circle). We found it convenient to combine these two plots into a single biplot (Gabriel, 1971). This allows to use two biplots (what we called a bibiplot) instead of four separate plots.

In ecology, reduced-rank regression has been long used for dimensionality reduction and data visualization, under the name of redundancy analysis (RDA) (Ter Braak, 1994; Ramette, 2007). This field uses biplots similar to the ones developed in this manuscript (Braak and Looman, 1994), sometimes combining both biplots into one figure. A recently suggested sparse RDA (Csala et al., 2017) has similar aim to our work; their method can be seen as a variant of sparse PLS.

### 3.3 Limitations and outlook

Following Friedman et al. (2010) and Chen and Huang (2012), we used group lasso that induces row-wise sparsity in **W**. This means that the same set of genes is selected for all RRR components. For *r* = 2, as used in this manuscript, the same set of genes influences the 1st and the 2nd component, which has both advantages and disadvantages. Our sparse RRR algorithm is easy to modify for the standard lasso case: using Σ∥*W*_*i·*_∥_1_ in Eq. (5) instead of Σ∥*W*_*i·*_∥_2_ would induce element-wise sparsity. In this case the loss in Eq. (7) can be minimized separately for each column of **W** (e.g. also using `glmnet`). Using this approach, different sets can be selected for different RRR components and the same value of *λ* can yield different number of selected genes for different components. However, when using relaxed elastic net and performing RRR again, all components will get nonzero contributions from all selected genes. Further work would be needed to formulate a relaxed version of the element-wise sparse RRR that would preserve element-wise sparsity. Empirically, we found that for the datasets considered here, the performance of the element-wise sparse RRR without relaxation was similar to the performance of the naive row-wise sparse RRR but worse than the performance of the relaxed row-wise sparse RRR.

One important caveat is that the list of selected genes should not be interpreted as definite. There are two reasons for that. First, the model performance (Figure 3) was unaffected in some range of parameters corresponding to selecting from ∼10 to ∼50 genes, meaning that the choice of regularization strength in this interval remains an analyst’s call. Second, even for fixed regularization parameters, a somewhat different set of genes may be selected every time the experiment is repeated. We used bootstrapping to directly estimate gene selection stability of sparse RRR, and found that the larger the sample size, the more stable the model was. There is an interplay between these two factors. Stronger *ℓ*_1_ regularization leads to a sparser model with less selection reliability. Weaker *ℓ*_1_ regularization leads to a less sparse model with higher selection reliability. We stress that such instability is an inherent feature of *all* sparse methods (Xu et al., 2011). We used bootstrapping to confirm that sparse CCA of Witten et al. had similar levels of selection instability.

In principle, it would be possible to generalize our regression framework to nonlinear mappings, using e.g. a neural network with a bottleneck instead of the low-rank linear mapping shown in Figure 1e. This can be an interesting direction for future research, but fitting such models would require sample sizes at least as high as in the M1 dataset.

In conclusion, we believe that sparse RRR can be a valuable tool for exploration and visualization of paired datasets. We expect that our method can be relevant beyond the scope of Patch-seq data. For example, spatial transcriptomics (Lein et al., 2017) combined with two-photon imaging may allow characterizing the transcriptome and physiology of individual cells in the intact tissue, yielding large multi-modal datasets. Similarly, other types of multi-omics data where single-cell or bulk transcriptomic data are combined with some other type of measurements (e.g. chemical, medical, or even behavioural), may benefit from interpretable visualization techniques such as the one introduced here.

## 4 Methods

### 4.1 Data preprocessing

#### L1 dataset

We used read count table from the original publication (Cadwell et al., 2016). In this dataset there are *n* = 51 interneurons (from 53 sequenced interneurons, 2 were excluded in the original publication as contaminated), *p* = 15 074 genes identified by the authors as detected, and *q* = 11 electrophysiological properties. We excluded all cells for which at least one electrophysiological property was not estimated, resulting in *n* = 44. We restricted the gene pool to the *p* = 3000 most variable genes, the same ones identified in the original publication. We used the expert classification of cells into two classes performed in the original publication for annotating cell types. Out of *n* = 44 cells, only 35 cells were classified unambiguously (score 1 or score 5 on the scale from 1 to 5); the remaining 9 cells received intermediate scores. When performing cross-validation with gene selection in the CV loop, we used the gene selection procedure from Kobak and Berens (2019).

#### S1 dataset

We used UMI counts table from the original publication (Fuzik et al., 2016). In this dataset there are *n* = 83 cells, *p* = 24 378 genes after excluding ERCC spike-ins, and *q* = 89 electrophysiological properties. Out of 83 sequenced cells, we were only able to match *n* = 80 to the electrophysiological data. We used only *q* = 80 electrophysiological properties for which the data were available for all these cells (the fact that *n* = *q* = 80 is coincidental). We selected *p* = 1 384 genes with average expression above 0.5 (before standardization) for the RRR analysis.

#### L4 dataset

We used read counts table from the original publication (Scala et al., 2019). In this dataset there are *n* = 110 Patch-seq neurons; we used the same *n* = 102 as in the original publication (after excluding low quality cells). We used the same *p* = 1000 genes selected in the original publication, and the same *q* = 13 electrophysiological properties.

#### M1 dataset

We used read counts table from the original publication (Scala et al., 2020). In this dataset there are *n* = 1320 Patch-seq neurons; we used the same *n* = 1213 as in the original publication (after excluding low quality cells). We used the same *p* = 1000 genes selected in the original publication, and the same *q* = 16 electrophysiological properties.

#### Preprocessing

For the full-length datasets (L1, L4, M1) we performed sequencing depth normalization by converting the counts to counter per million (CPM). For the UMI-based dataset (S1) we divided the values for each cell by the cell sum over all genes (sequencing depth) and multiplying the result by the median sequencing depth size across all cells. In both cases we then log-transformed the data using log2(*x* + 1) transformation. Finally, we standardized all gene expression values and all electrophysiological properties to zero mean and unit variance.

### 4.2 Data availability

All datasets were either downloaded from the original publications or provided by the authors. All of them can be found at https://github.com/berenslab/patch-seq-rrr. Our full analysis code in Python is also available there.

## 5 Appendix. Procrustes problem

Given **A**, the Procrustes problem is to maximize tr(**AV**^⊤^) subject to **V**^⊤^**V** = **I** (Gower and Dijksterhuis, 2004). Let us denote by 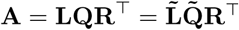 the thin and the full SVD of **A**. Now we have:

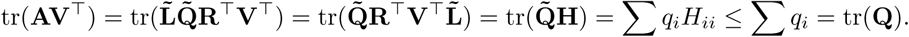

Here 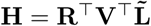 is a matrix with orthonormal rows as can be verified directly, and so it must have all its elements not larger than one. It follows that the whole trace is not larger than the sum of singular values of **A**. Using **V** = **LR**^⊤^ yields exactly this value of the trace, hence it is the optimum.

## Authors’ contributions

PB and DK conceptualized the project, DK developed the statistical method and wrote the software, DK, YB, and MW performed computational experiments, FS performed Patch-seq experiments under the supervision of AT and helped analyzing the data, PB supervised the project. DK and PB wrote the paper with input from all authors.

## Acknowledgements

We thank Rickard Sandberg, Cathryn Cadwell, and Jiaolong Xiang for discussions, Cathryn Cadwell, Janos Fuzik and their co-authors for making their data available and Shreejoy Tripathy for comments and help with data processing. This work was funded by the German Ministry of Education and Research (FKZ 01GQ1601), the German Research Foundation (EXC 2064 project number 390727645, BE5601/4-1) and the National Institute Of Mental Health of the National Institutes of Health under Award Number U19MH114830. The content is solely the responsibility of the authors and does not necessarily represent the official views of the National Institutes of Health.

## Footnotes

1 In some related previous work (González et al., 2008, 2009), cross-validated correlations were computed by pooling test set points across all cross-validation splits. We observed that this procedure can sometimes yield biased results; we compute test-set correlation within each test set, and then average across CV splits.

